# Differential gene expression indicates involvement of F-box proteins and E3 ligases in sexual *versus* apomictic germline specification in *Boechera*

**DOI:** 10.1101/403915

**Authors:** Luise Zuehl, David Ibberson, Anja Schmidt

## Abstract

Germline specification is the first step during sexual and apomictic plant reproduction. This takes place in a specialized domain of the reproductive flower tissues, the nucellus of the ovule. In each case, a sporophytic cell is determined to initiate germline development. These cells, the megaspore mother cell (MMC) or apomictic initial cell (AIC) in sexual plants and apomicts, respectively, differ in their developmental fate. While the MMC undergoes meiosis, the AIC aborts or omits meiosis to form the female gametophyte. Although these distinct developmental processes have long been described, little is known about the gene regulatory basis involved.

To elucidate gene regulatory networks underlying sexual and apomictic germline specification, we conducted tissue-specific transcriptional profiling using laser-assisted microdissection and RNA-Seq. We compared the transcriptomes of the nucellar tissues harbouring the MMC or AIC between different accessions of *Boechera*. The six accessions we used represented four species and two ploidy levels, allowing us to distinguish between differences in gene expression caused by the genetic background or the reproductive mode.

Comparative data analysis revealed widely overlapping gene expression patterns in apomictic *versus* sexual *Boechera* accessions. Nevertheless, 45 significantly differentially expressed genes were identified, which potentially play a role for determination of sexual *versus* apomictic reproductive mode. Interestingly, based on annotations, these include F-box proteins and E3 ligases that might relate to genes previously described as regulators important for sexual or apomictic reproduction. Thus, our findings provide new insight into the transcriptional basis of sexual and apomictic germline specification.

**One sentence summary**

A comprehensive tissue type-specific transcriptional analysis using laser-assisted microdissection combined with RNA-Seq identifies 45 genes consistently differentially expressed during germline specification in different sexual *versus* apomictic *Boechera* accessions, indicating roles of protein degradation related to cell cycle, transcriptional and post-transcriptional regulatory processes, and stress response for apomixis.

## INTRODUCTION

The acquisition of reproductive or germline fate is a key step in the life cycle of higher plants, as it mediates the transition from the sporophytic to the gametophytic generation. The formation of the female germline initiates with the selection of a single sporophytic cell of the nucellus for reproductive fate, the megaspore mother cell (MMC). In sexually reproducing plants, the MMC undergoes meiosis to generate the functional megaspore (FMS) as the first cell of the gametophytic generation. During gametogenesis, the mature female gametophyte is formed from the FMS by three rounds of mitosis followed by cellularization. The mature gametophyte consists of the two female gametes, the egg- and the central cell, and accessory synergid cells and antipodals (Sprunck and Gross-Hardt, 2011). Double fertilization of the egg- and central cell, giving rise to embryo and endosperm, respectively, completes the life cycle by the start of a new sporophytic generation.

Apart from sexual reproduction through seeds, more than 400 angiosperm species of approx. 40 families, produce seeds asexually in a process called gametophytic apomixis (Asker and Jerling, 1992; Carman, 1995; Spillane et al., 2001). Gametophytic apomixis (hereafter referred to as apomixis) comprises three consecutive developmental processes during reproduction differing from the sexual pathway. It begins with the formation of an unreduced female gametophyte (apomeiosis), which subsequently enables fertilization-independent embryogenesis (parthenogenesis) and fertilization-dependent (pseudogamous) or autonomous development of functional endosperm (Nogler, 1984; Asker and Jerling, 1992). In general, two major types of apomixis, diplospory and apospory, can be distinguished based on how the female germline is established (Nogler, 1984; Asker and Jerling, 1992; Koltunow, 1993; Bicknell and Koltunow, 2004). In diplospory, a single apomictic initial cell (AIC) is specified instead of the MMC at the same position in the ovule. The AIC either omits or enters meiosis, which however aborts, leading to the formation of an unreduced FMS. In contrast, aposporous development is characterised by an aposporous initial cell (also AIC), which specifies adjacent to a sexual MMC. This cell fully circumvents meiosis to differentiate into an unreduced FMS. In each case, the FMS further develops into a mature gametophyte as in sexually reproducing plants, however gametes typically remain genetically identical to the mother plant. Followed by parthenogenesis of the egg cell, apomictic reproduction thus leads to the formation of clonal offspring.

As it is foreseen that complex genotypes from hybrid crops can be fixed over subsequent generations by harnessing apomixis, it holds great promises for uses in agriculture. Hence, the understanding of germline specification in both sexuals and apomicts is of special scientific interest as such, but also regarding the potential of apomixis for agricultural application. Despite increasing interest in the developmental processes leading to apomictic reproduction, knowledge about the underlying genetic and transcriptional basis is still scarce.

A widely accepted hypothesis states that apomixis results from asynchronous expression of genes of the sexual pathway (Carman, 1997). This is likely due to the hybrid origin or polyploid nature of most apomicts (Bicknell and Koltunow, 2004; Lovell et al., 2013). Meanwhile, a strong body of evidence implies that apomixis evolved from a deregulation of the sexual reproductive pathway several times independently (Koltunow, 1993; Grimanelli et al., 2001; Grossniklaus et al., 2001; Koltunow and Grossniklaus, 2003; Tucker et al., 2003; Sharbel et al., 2009; Sharbel et al., 2010).

An increasing number of genetic and molecular studies in sexual model species such as *Arabidopsis thaliana*, *Zea mays* or *Oryza sativa*, provides strong evidence that complex regulatory networks are required for megasporogenesis and germline development (Schmidt et al., 2015; Nakajima, 2018). In contrast to aposporous apomicts, where the AIC typically forms adjacent to the sexual MMC, the formation of only one germline in each ovule is tightly controlled in sexual species (Albertini et al., 2005; Albrecht et al., 2005; Colcombet et al., 2005; Olmedo-Monfil et al., 2010; Schmidt et al., 2011; Singh et al., 2011; Wang et al., 2012a; Singh et al., 2017; Zhao et al., 2017b; Cao et al., 2018). Interestingly, mutations in a number of genes, important for regulation of sexual germline development, lead to aposporous- or diplosporous-like phenotypes in sexual model species (Barcaccia and Albertini, 2013; Hand and Koltunow, 2014; Schmidt et al., 2015). These include such regulatory components, which are involved in restricting reproductive fate (Hand and Koltunow, 2014; Schmidt et al., 2015). Developmental alterations resembling apospory are described for mutations disrupting members of epigenetic regulatory and small RNA pathways in *A. thaliana* and maize, e.g. *A. thaliana ARGONAUTE9* (*AGO9*), *RNA-DEPENDENT RNA POLYMERASE6* (*RDR6*) and *SUPPRESSOR OF GENE SILENCING3* (*SGS3*) (Ravi et al., 2008; Garcia-Aguilar et al., 2010; Olmedo-Monfil et al., 2010; Singh et al., 2017). In contrast, phenotypes resembling diplospory are induced in mutant lines of *AGO104* in maize, a homologue of *A. thaliana AGO9*, in addition to *DYAD* and other mutants in core meiotic genes (Ravi et al., 2008; d‘Erfurth et al., 2009; Singh et al., 2011). However, if alterations in these genes and pathways are underlying reproductive development in natural apomicts remains to be elucidated.

To study apomictic reproduction, the genus *Boechera* Á. Löve & D. Löve is increasingly used as a model (Sharbel et al., 2009; Aliyu et al., 2010; Sharbel et al., 2010; Aliyu et al., 2013; Corral et al., 2013; Lovell et al., 2013; Schmidt et al., 2014; Mau et al., 2015; Rojek et al., 2018). While apomixis is almost exclusively restricted to polyploids in most species, it occurs at low ploidy levels including diploid and triploid throughout the genus (Windham and Al-Shehbaz, 2007). In *Boechera,* both apospory and diplospory have been described (Aliyu et al., 2010; Mateo de Arias, 2015). However, the underlying molecular processes are not fully understood.

Based on the hypothesis of apomixis being derived from a deregulation of the sexual pathway, transcriptional profiling has previously been used to identify differential regulation during sexual and apomictic reproduction. In particular for transcriptome analyses of the female germline, cell or tissue type-specific investigations bear great potential, as otherwise important genes might be overlooked due to overabundance of sporophytic tissues in the sample (Wuest et al., 2010; Schmid et al., 2012; Schmidt et al., 2012; Florez Rueda et al., 2016). Previous studies in *Boechera* compare entire ovules of sexual and apomictic plants, while cell type-specific investigations have been reported comparing sexual MMC of *A. thaliana* and apomictic AIC of *B. gunnisoniana* (Sharbel et al., 2009; Sharbel et al., 2010; Schmidt et al., 2014). This revealed several hundreds of genes to be differentially expressed in apomictic *versus* sexual germlines, which are involved in cell cycle control, epigenetic, transcriptional and hormonal regulation (Schmidt et al., 2014).

To narrow down genes and pathways of putative importance for apomixis and to minimize confounding effects of ploidy or species differences, we now performed a comprehensive investigation of two sexual and four apomictic *Boechera* accessions. We used laser-assisted microdissection (LAM) combined with RNA-Seq to perform a comparative transcriptome analysis of nucellus tissues harbouring the MMC or AIC. This allows to study specification of the germline in a tissue-specific manner. Moreover, any relevant signalling processes involving the cells of the sporophytic nucellus can simultaneously be detected. Thereby, 45 genes consistently differentially regulated between sexual and apomictic accessions were identified, pointing to the involvement of genes and pathways previously described as regulators for germline development, i.e. cell cycle regulation and stress response.

## RESULTS

### Tissue-specific transcriptome profiling of sexual and apomictic nucelli

To gain new insights into the gene regulatory networks underlying germline specification during sexual and apomictic reproduction tissue-specific RNA-Seq was performed. The study design comprised four obligatory apomictic and two sexually reproducing accessions of the genus *Boechera* (Table 1) (Schranz et al., 2005; Aliyu et al., 2010; Mau et al., 2015; Lee et al., 2017). They were selected based on reproductive mode in the first place, but also to represent four different species and two ploidy levels (Table 1). This allows to distinguish differential expression based on reproductive mode from effects of ploidy or genetic background of the different species.

**Table 1.** *Boechera* accessions included in this study. All accessions were previously described as indicated. Accessions were selected to represent different reproductive modes (sexual, apomictic), ploidies (2n, 3n), and four different species.

| **Species** | **Accession no.** | **Reproductive mode** | **Ploidy** | **Reference** |
| --- | --- | --- | --- | --- |
| *Boechera divaricarpa* | ES517 | apomictic | 2n | Aliyu et al. (2010) |
| *Boechera pallidifolia* | B12-1578 | apomictic | 2n | Mau et al. (2015) |
| *Boechera pallidifolia* | B12-1599 | apomictic | 3n | Mau et al. (2015) |
| *Boechera williamsii* | B12-1524 | apomictic | 2n | Mau et al. (2015) |
| *Boechera williamsii* | B12-558 | sexual | 2n | Mau et al. (2015) |
| *Boechera stricta* | LTM | sexual | 2n | Schranz et al. (2005), Lee et al. (2017) |

LAM was used to isolate nucelli from developing ovules harbouring the MMC or AIC (Figure 1). Nucellus tissues were precisely dissected from dry sections of young floral buds with only about 2 μm cutting width of laser beam (Figure 1B). However, minor cross-contamination with surrounding sporophytic tissue could not be avoided completely, due to the small size of nucelli and technical constrains. In order to identify genes consistently differentially regulated between all sexual and all apomictic accessions included in the analysis, a total of 9 samples was prepared including biological replicates for one sexual and two apomictic accessions (LTM, ES517, B12-1578; Table S1).

**Figure 1.**
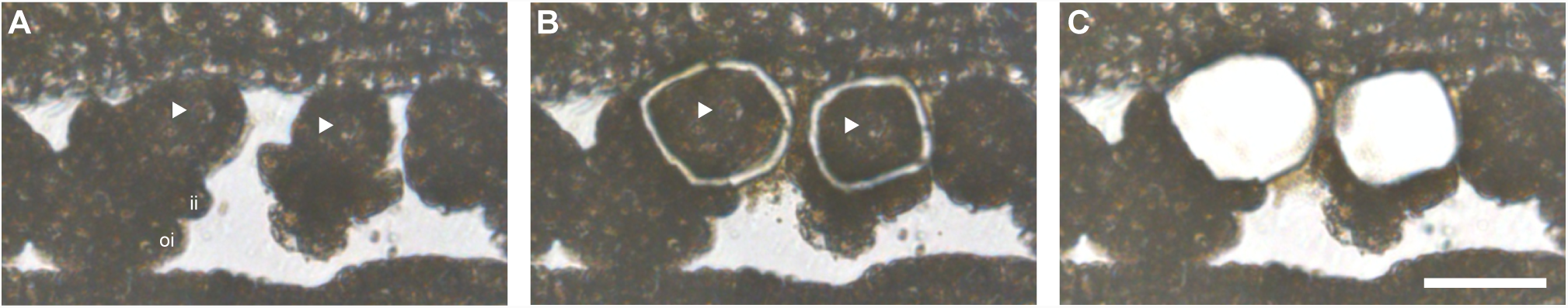
Precise collection of nucelli using laser-assisted microdissection. Nucelli harbouring AIC in ovary of apomictic *B. pallidifolia* (B12-1578, 2n) **(A)** before, **(B)** after LAM and **(C)** after sample collection. Nuclei of AIC are marked with arrow heads and inner (ii) - and outer (oi) integuments are labelled, accordingly. Scale bar = 30 μm.

For transcriptional profiling, the RNA was extracted, amplified, and subjected to library preparation (Figure S1). Subsequently, RNA-Seq was performed using the single-end 75 bp protocol with 19 additional cycles on one lane of the Illumina NextSeq 500 platform. Individual libraries comprised 28,396,884 – 99,921,406 reads of overall good quality (63 million reads on average; Table S1).

After quality controls and trimming, reads were mapped to the *B. stricta* reference genome with STAR (Dobin et al., 2013; Lee et al., 2017). On average about 98% of the reads could be mapped per library (Table S1). Unique reads constituted 95% of total reads on average, and gene bodies were covered homogenously by mapped reads (Table S1, Figures S2). Further, 85% of total reads mapped to exonic regions, when counted with featureCounts (Table S1) (Liao et al., 2014). Taken together, quality controls performed during processing of RNA-Seq reads, general composition of read sequences and library statistics demonstrated overall good quality of RNA-Seq data obtained (Table S1, Figures S2).

### Global gene expression in apomictic and sexual *Boechera* nucelli widely overlapped

In this study, samples from nucellus tissues at similar developmental stages isolated from closely related sexual and apomictic *Boechera* accessions were profiled. This provides a good basis to identify genes consistently transcribed, and thus likely of general importance independent of the reproductive mode. Importantly however, genes differentially regulated in apomictic and sexual nucelli can be identified, which are supposedly relevant for determination and progression of either sexual or for apomictic germline development.

All together, we identified 24,197 genes to be expressed (≥ 10 normalized read counts) in at least one of the samples analysed, thus representing 88% of 27,416 genes in *B. stricta* (Lee et al., 2017). Of those, 16,221 genes were expressed either in all of the samples analysed from apomicts, or all from sexual accessions, or both (Figure 2). According to our expectations for profiling similar tissue types of closely related accessions, consistent expression in all samples was detected for 10,526 genes, representing 38% of the genes in the genome (Figure 2). Expression of 710 genes was shared only in the samples from the sexual accessions but not in the apomicts, while exclusive expression in all apomictic samples was observed for 4,985 genes (Figure 2). Among all samples from apomicts however, expression of 15,511 genes was shared (Figure S3A). And in contrast, only 11,236 genes were found to be expressed in all sexual samples (Figure S3B). Despite similar regulation for a large number of genes, also variation of expression between species or accessions of the same reproductive mode was observed (Figure S3).

**Figure 2.**
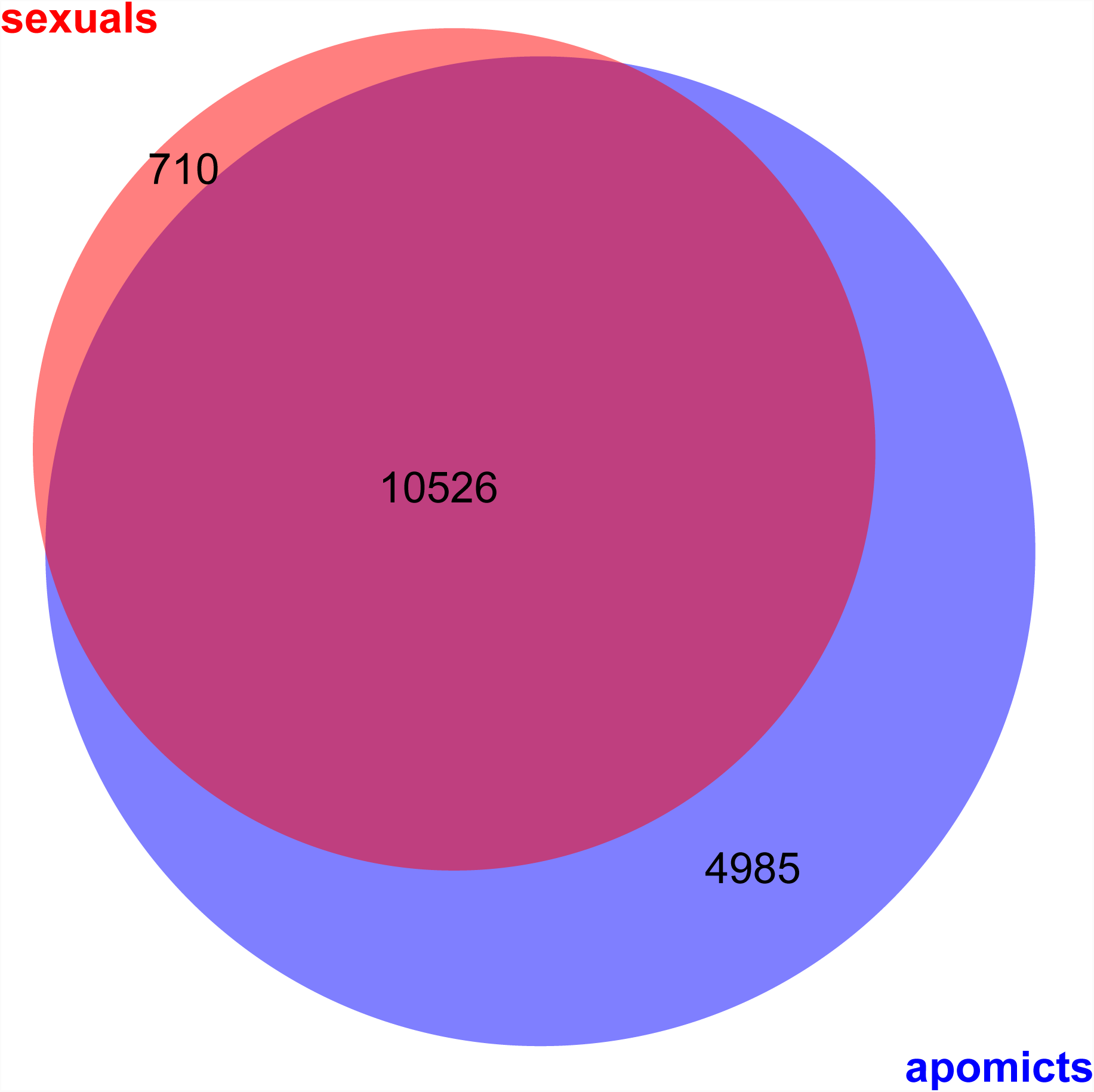
Comparison of expressed genes in nucelli of different reproductive modes. Venn diagram represents overlap of genes expressed commonly in either all samples from apomictic or all from sexual accessions (≥ 10 normalized read counts).

In summary, comparative analyses revealed widely overlapping gene expression patterns in nucelli of apomictic and sexually reproducing *Boechera* accessions. Nevertheless, the comparisons already indicated distinct regulation of a subset of genes according to the reproductive mode.

### Statistical analysis reveals genes consistently differentially expressed between apomictic and sexual nucelli

Despite the large number of genes consistently expressed in the nucelli tissues of all accessions independent of the reproductive mode, differential expression for genes important for determination of the reproductive mode or developmental processes specific to either reproductive mode is expected. To identify genes of potential relevance for determination of the reproductive mode, statistical analysis of the differential gene expression between apomictic and sexual nucelli samples was applied using edgeR (Robinson et al., 2010).

First, we compared the transcriptomes of all apomictic against all sexual samples, treating samples of same reproductive mode as biological replicates. With this approach 43 genes were identified to be significantly differentially expressed between samples originating from apomictic and those from sexual nucelli (*p* values ≤ 0.05 after Benjamini-Hochberg adjustment of FDR; Figure 3, Table S2, S3). Of these differentially expressed genes (DEGs) 28 were significantly upregulated in apomictic and 18 in sexual samples (Figure 3, Table S2, S3). Hierarchical clustering and heatmap representation of their expression levels demonstrated a clear distinction between reproductive modes and close relation between biological replicates (Figure 3). Moreover, expression patterns of samples from *B. pallidifolia* accessions (B12-1578 and B12-1599) clustered together, as well as those of apomictic *B. divaricarpa* (ES517) with the apomictic *B. williamsii* (B12-1524) accessions (Figure 3).

**Figure 3.**
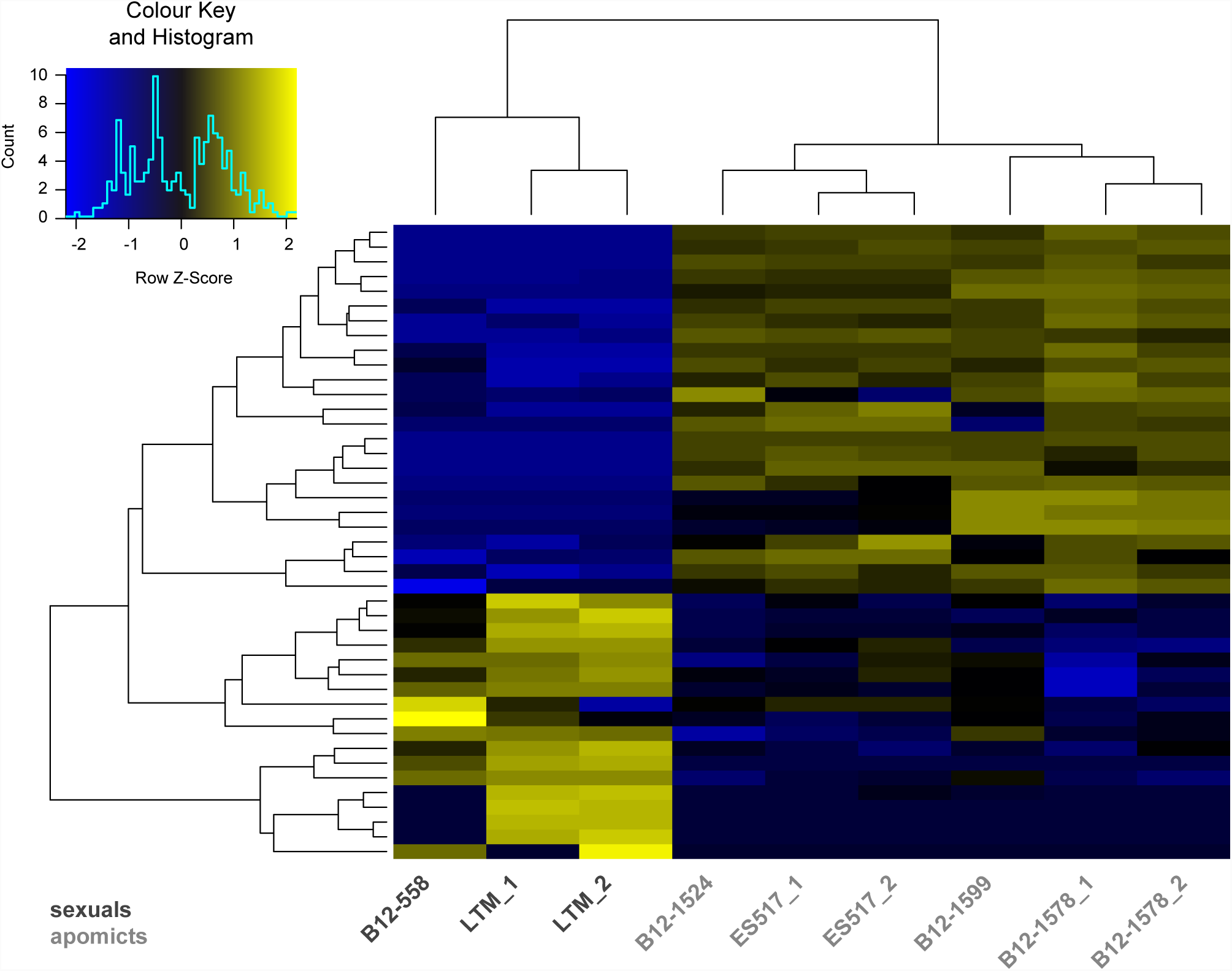
Heatmap of log2 transformed read counts of differentially expressed genes (DEGs). Expression levels of 43 DEGs in all sexual as compared to all apomictic samples. The heatmap is based on TMM normalized and log2 transformed read counts. The hierarchical clustering of samples and genes was based on Euclidean distance and hierarchical agglomerative clustering. Colours of heatmap are scaled per row with blue denoting low and yellow high expression.

In a second approach, ploidy and genetic backgrounds of analysed accessions were taken into account. To this end, DEGs were first identified by individual pairwise comparisons of all samples from sexual against all samples from apomictic accessions (*p* values ≤ 0.05 after Benjamini-Hochberg adjustment of FDR; Table S3). Relevant candidate genes for germline specification were narrowed down by taking the intersection of all DEGs identified in individual pairwise comparisons. This resulted in the identification of 7 common DEGs, which were all higher expressed in apomictic than in sexual accessions (Figure S4). Five of these have consistently been identified in both analyses (Table 2, Table S4).

**Table 2.** Differentially expressed genes (DEGs) between all samples originating from apomictic and all from sexual nucelli. **(A)** DEGs upregulated either in apomictic accessions or **(B)** in sexual accessions. In total 43 are listed with information on *Boechera stricta* gene locus, GO terms and *A*. *thaliana* homologues based on genome annotations by Lee et al. (2017). DEGs are ordered according to Table S2 Sheet 1 and those identified consistently in both analyses are highlighted in grey. NA = no information available.

| Gene locus | GO terms | *A. thaliana*<br>gene locus | *A. thaliana*<br>gene name | *A. thaliana* gene/ protein description |
| --- | --- | --- | --- | --- |
| **(A) DEGs upregulated n apomictic accessions:** |  |  |  |  |
| Bostr.26527s0134 | GO:0030246, GO:0005975,<br>GO:0004553 | AT5G20710 | BGAL7 | Beta-galactosidase 7 |
| Bostr.7867s0569 | GO:0046872 | AT4G27470 | ATRMA3, RMA3 | RING membrane-anchor 3 |
| Bostr.25375s0042 | NA | AT1G64290 | NA | F-box protein-related |
| Bostr.15697s0319 | NA | AT1G30790 | NA | F-box and associated interaction domains-containing protein |
| Bostr.15697s0321 | NA | AT1G32660 | NA | F-box and associated interaction domains-containing protein |
| Bostr.13129s0275 | NA |  | NA | NA |
| Bostr.29044s0001 | NA | AT5G36710 | NA | NA |
| Bostr.3288s0214 | GO:0033177, GO:0015991,<br>GO:0015078 | AT1G19910 | ATVHA-C2, AVA-2PE, AVA-P2 | ATPase, F0/V0 complex, subunit C protein |
| Bostr.2983s0066 | GO:0005515 | AT4G05310 | NA | Ubiquitin-like superfamily protein |
| Bostr.15697s0461 | GO:0046872 | AT1G31480 | SGR2 | Shoot gravitropism 2 (SGR2) |
| Bostr.0568s0078 | GO:0008270 | AT5G17890 | CHS3, DAR4 | DA1-related protein 4 |
| Bostr.0697s0107 | NA | AT4G10400 | NA | F-box/RNI-like/FBD-like domains-containing protein |
| Bostr.26833s0971 | NA | NA | NA | NA |
| Bostr.19424s0775 | NA | NA | NA | NA |
| Bostr.10058s0023 | GO:0008168 | AT4G00740 | NA | S-adenosyl-L-methionine-dependent methyltransferases<br>superfamily protein |
| Bostr.13158s0243 | GO:0055114, GO:0009396,<br>GO:0004488, GO:0003824 | AT4G00620 | NA | Amino acid dehydrogenase family protein |
| Bostr.26959s0357 | GO:0055114, GO:0009396,<br>GO:0004488, GO:0003824 | AT4G00620 | NA | Amino acid dehydrogenase family protein |
| Bostr.7305s0030 | NA | AT5G52390 | NA | PAR1 protein |
| Bostr.18351s0192 | GO:0005515 | AT2G17310 | SON1 | F-box and associated interaction domains-containing protein |
| Bostr.3279s0010 | GO:0010468, GO:0005777 | AT4G30760 |  | Putative endonuclease or glycosyl hydrolase |
| Bostr.26833s0821 | NA | AT5G54510 | DFL1, GH3.6 | Auxin-responsive GH3 family protein |

**Table 2 continued.**
| Gene locus | GO terms | *A. thaliana*<br>gene locus | *A. thaliana*<br>gene name | *A. thaliana* gene/ protein description |
| --- | --- | --- | --- | --- |
| Bostr.5022s0132 | NA | AT2G21140 | ATPRP2, PRP2 | Proline-rich protein 2 |
| Bostr.25993s0534 | NA | AT2G42480 | NA | TRAF-like family protein |
| Bostr.13158s0246 | GO:0055114, GO:0009396,<br>GO:0004488, GO:0003824 | AT4G00620 | NA | Amino acid dehydrogenase family protein |
| Bostr.13671s0276 | GO:0000062 | AT1G31812 | ACBP, ACBP6 | Acyl-CoA-binding protein 6 |
| **(B) DEGs upregulated in sexual accessions:** |  |  |  |  |
| Bostr.13129s0114 | GO:0005515, GO:0003677 | AT5G08430 | NA | SWIB/MDM2 domain; Plus-3; GYF |
| Bostr.26959s0127 | NA | AT1G67800 | NA | Copine (Calcium-dependent phospholipid-binding protein) family |
| Bostr.7867s0507 | GO:0005975, GO:0004553 | AT4G26830 | NA | O-Glycosyl hydrolases family 17 protein |
| Bostr.12396s0001 | GO:0055114, GO:0051287,<br>GO:0016616 | AT1G26570 | ATUGD1, UGD1 | UDP-glucose dehydrogenase 1 |
| Bostr.7867s1023 | GO:0003677 | AT4G31680 | NA | Transcriptional factor B3 family protein |
| Bostr.18351s0250 | GO:0007165, GO:0000155,<br>GO:0000160 | AT2G17820 | AHK1, ATHK1, HK1 | Histidine kinase 1 |
| Bostr.7867s1594 | GO:0055114, GO:0020037,<br>GO:0016705, GO:0005506 | AT4G37330 | CYP81D4 | Cytochrome P450, family 81, subfamily D, polypeptide 4 |
| Bostr.0697s0084 | GO:0055085, GO:0016021 | AT3G54700 | PHT1;7 | Phosphate transporter 1;7 |
| Bostr.26833s0862 | GO:0016758, GO:0008152 | AT3G21790 | NA | UDP-Glycosyltransferase superfamily protein |
| Bostr.15774s0342 | GO:0055114, GO:0016614,<br>GO:0050660 | AT5G51950 | NA | Glucose-methanol-choline (GMC) oxidoreductase family protein |
| Bostr.2021s0098 | NA | NA | NA | NA |
| Bostr.29223s0097 | NA | AT1G64230 | UBC28 | Ubiquitin-conjugating enzyme 28 |
| Bostr.2618s0047 | GO:0005975, GO:0004553 | AT5G17500 | NA | Lycosyl hydrolase superfamily protein |
| Bostr.15697s0040 | NA | NA | NA | NA |
| Bostr.7867s0172 | NA | AT1G25290 | ATRBL10, RBL10 | RHOMBOID-like protein 10 |
| Bostr.26833s0734 | NA | AT5G55240 | ATPXG2 | ARABIDOPSIS THALIANA PEROXYGENASE 2 |
| Bostr.25993s0569 | NA | AT3G53750 | ACT3 | Actin 3 |
| Bostr.3751s0034 | NA | AT5G40680 | NA | Galactose oxidase/kelch repeat superfamily protein |

Despite a large overlap of global gene expression in apomictic and sexual nucelli, statistic data analysis identified 45 genes to be significantly differentially expressed in all sexual *versus* all apomictic samples generated in this study (Table S3). Due to the comparison of closely related *Boechera* accessions, reducing the influence of varying genetic backgrounds, this set of DEGs contains promising new candidates of putative importance for the distinct germline specification in apomictic *versus* sexually reproducing *Boechera.*

### Annotations of DEGs indicate putative roles in protein degradation, transcriptional regulation and stress response

Since *Boechera* is a genus used as a model only relatively recently, knowledge about the specific gene functions in *Boechera* is scarce. However, the close relation of *Boechera* to *A. thaliana* and genome annotations of *B. stricta* including information on corresponding *A. thaliana* homologues to *B. stricta* genes are a useful basis for investigations on *Boechera* and comparative analyses (Schmidt et al., 2014; Lee et al., 2017).

In numbers, 21,890 *A. thaliana* genes are annotated as homologues of *B. stricta* genes, thus covering approx. 80% of the 27,416 genes in the *B. stricta* genome (Lee et al., 2017). Accordingly, corresponding *A. thaliana* are annotated for the majority of DEGs identified. Based on this, the DEGs could be attributed to different functional categories: (1) protein degradation, (2) transcriptional and post-transcriptional regulation of gene expression, (3) redox processes and stress response, and (4) phytohormone and cell signalling (Tables 2, Table S4).

The first group of genes related to protein degradation comprised especially genes coding for proteins with F-box and associated interaction domains (e.g. cyclin-like and Skp2-like domains), a F-box/RNI-like superfamily protein or a protein related to F-box proteins. These genes were mostly upregulated in apomictic nucelli (Table 2A, Table S4). Additionally, two E3 ligases were identified to be differentially expressed. Bostr.7867s0569, the homologue of the E3 ligase *RING MEMBRANE-ANCHOR 3 (ATRMA3)* was consistently identified in both comparisons (Tables 2A, Table S4). And a gene encoding a Tumor Necrosis Factor Receptor (TNF-R) Associated Factors like (TRAF-like) family protein, Bostr.25993s0534, was found to be significantly upregulated in apomictic nucelli (Tables 2A, Tables S4). Apart from this, two genes related to ubiquitination were represented in the set of DEGs: the homologues of the *UBIQUITIN-CONJUGATING ENZYME 28 (UBC28),* and of an ubiquitin-like superfamily protein, Bostr.29223s0097 and Bostr.2983s0066, respectively (Table 2, Table S4).

Secondly, two DEGs related to transcriptional and post-transcriptional regulation of gene expression were identified. A putative endonuclease, Bostr.3279s0010, was significantly upregulated in apomictic nucelli in both analyses, whereas a B3 family transcription factor, Bostr.7867s1023, was identified to be significantly lower expressed in apomictic than sexual nucelli (Table 2A, Tables S4).

The third group of DEGs included genes with *A. thaliana* homologues related to redox processes and stress responses. Most prominently, a gene encoding a vacuolar ATPase subunit protein, Bostr.3288s0214, was found to be significantly higher expressed in apomicts in both comparisons, which has previously been described to be related to abiotic stress response (Kreps et al., 2002) (Tables 2A, Tables S4). In contrast, several candidate genes of this third group were significantly less expressed in apomictic than in sexual nucelli. This subset was comprised of the homologue of *ARABIDOPSIS THALIANA PEROXYGENASE 2 (ATPXG2)*, Bostr.26833s0734, a gene coding for a glucose-methanol-choline (GMC) oxidoreductase family protein, Bostr.15774s0342 and another oxidoreductase, i.e. a member of the cytochrome P450 superfamily *CYP81D4,* Bostr.7867s1594 (Table 2B, Tables S4).

In addition, a fourth group of genes was identified to be differentially expressed, of which *A. thaliana* homologues are involved in phytohormone-mediated cell signalling, or cell signalling *per se*. A gene coding for an auxin-responsive GRETCHEN HAGEN 3 (GH3) family protein (GH3.6), Bostr.26833s0821, and the homologue of *A. thaliana* nonethylene receptor HISTIDIN KINASE 1, Bostr.18351s0250, which is a positive regulator in ABA signal transduction and drought response (Tran et al., 2007) (Table 2, Tables S4).

Interestingly, two so far undescribed genes, Bostr.13129s0275 and Bostr.19424s0775, that lack any annotation on the reference genome, were consistently detected in both approaches, apart from the homologue of *ATRMA3*, the putative endonuclease and the gene encoding the V-ATPase subunit protein (Table 2A, Tables S4).

In summary, the dataset indicates consistent differential regulation of expression between sexual and apomictic germlines for a rather small number of genes. The identified differences mainly point towards involvement of poly-ubiquitination mediated protein degradation, transcriptional and post-transcriptional regulatory mechanisms, regulation by plant hormones, signal transduction and stress response. In addition, few genes with unknown functions sharing no significant homology to *A. thaliana* genes are identified among the set of DEGs.

### Comparisons to transcriptomes of *A. thaliana* MMC and *B. gunnisoniana* AIC indicates overall accuracy of dataset

Previously, cell type- and tissue type-specific transcriptome analyses have been described for the *A. thaliana* MMC and surrounding tissues of the nucellus isolated separately, and the AIC of the triploid apomict *B. gunnisoniana* (Schmidt et al., 2011; Schmidt et al., 2014). To identify genes sharing expression upon sexual or upon apomictic germline formation transcriptional profiles were compared. However, comparisons were restricted to commonly annotated *A. thaliana* homologues because available expression data are either generated with the Affymetrix ATH1 microarray for *A. thaliana* (Schmidt et al., 2011), or reads were mapped and annotated using a reference transcriptome from *B. gunnisoniana* in case of RNA-Seq data (Schmidt et al., 2014).

First, overlapping gene expression between MMC or sporophytic nucellar tissue without MMC of *A. thaliana* and transcriptomes of nucelli from all analysed sexual *Boechera* accessions (B12.558, LTM_1, LTM_2) was determined. Genes were defined to be expressed if they had either ≥ 10 normalized read counts or present calls in ≥ 3 of 4 microarray replicates for *A. thaliana* data (Schmidt et al., 2011). Overlapping expression in all samples of sexual *Boechera* nucelli tissues as compared to the transcriptomes of the *A. thaliana* MMC or nucellus tissues was identified for 5,139 of 6,650 (77%) or for 7267 of 10,081 (72%) genes, respectively (Figure 4A). Besides, approx. 91% (10,940) of 11,967 genes expressed (≥ 10 reads) in AIC of *B. gunnisoniana* were also identified to be active (≥ 10 reads) in the triploid *B. pallidifolia* accession analysed in this study (Figure 4B) (Schmidt et al., 2014).

**Figure 4.**
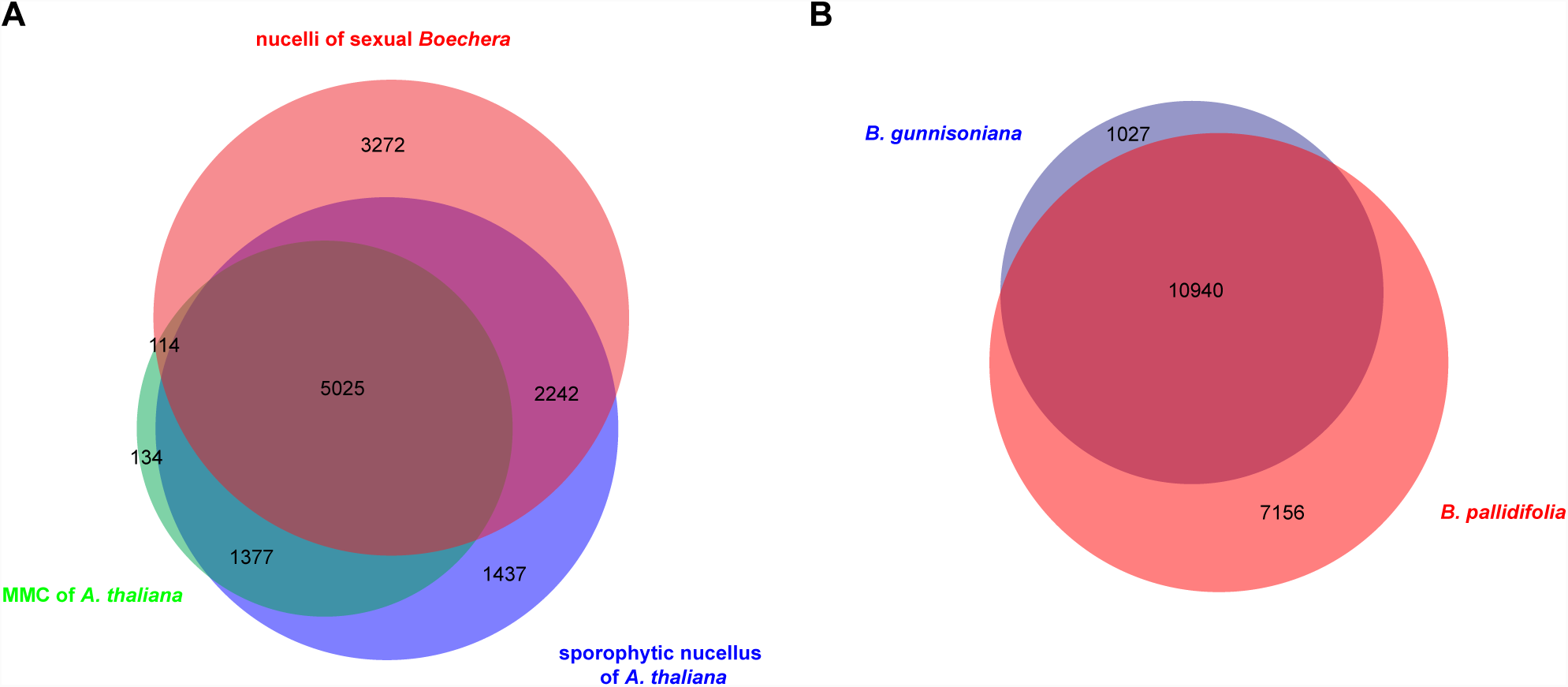
Common gene expression in female reproductive tissues of *A. thaliana* and *Boechera* accessions. **(A)** Venn diagram represents intersections of genes expressed in isolated MMC or sporophytic nucellar tissue of *A. thaliana* (present calls in ≥ 3 of 4 of microarray replicates (Schmidt et al., 2011)) and nucelli of all herein analysed samples of sexual *Boechera* accessions (LTM, B12. 558). **(B)** Venn diagram represents overlap of expressed genes (≥ 10 reads) in AIC of triploid *B. gunnisoniana* (Schmidt et al., 2014) and nucelli of *B. pallidifolia* (B12.1599, this study). Each comparison was based annotated *A. thaliana* homologues of corresponding *Boechera* genes; in **(A)** only those *A. thaliana* genes were considered which are represented on the Affymetrix ATH1 microarray.

In conclusion, high overlap of gene expression revealed by comparison of the nucelli transcriptomes to the transcriptomes of MMC and sporophytic nucelli in *A. thaliana* and AIC in *B. gunnisoniana* indicates an overall high accuracy of the dataset presented in this study.

## DISCUSSION

### Genes involved in cell cycle regulation, transcriptional control and stress response are differentially regulated during sexual and apomictic germline specification

The identification of genes of putative relevance to determine the reproductive mode or to sustain either sexual or apomictic development remains challenging, due to the small size of the reproductive tissues. Previously, a comparison of gene activity between triploid *B. gunnisoniana* and sexual *A. thaliana* has identified about 900 genes with evidence of expression in the AIC unlike in the MMC (Schmidt et al., 2014). While this study allowed important insights in the gene regulatory processes in the respective cell types, based on the number of genes it remained difficult to determine those important for apomictic in contrast to sexual development.

In the presented study, nucelli of same or closely related apomictic and sexual *Boechera* accessions were compared. This study design, also including accessions of two ploidy levels, allowed comparisons largely reducing disturbing effects of ploidy or species differences. Assuming that genes responsible for apomixis are the same in all *Boechera* accessions compared. By this, the number of genes identified with putative roles for determination of the reproductive mode could be narrowed down to 45. Based on gene annotations, they can be functionally attributed to different regulatory mechanisms, including transcriptional and post-transcriptional regulatory processes, protein degradation, cell signalling, and stress response. This is in agreement with previous findings and hypotheses (Schmidt et al., 2014; Mateo de Arias, 2015; Schmidt et al., 2015; Tang et al., 2017). Genes identified to be consistently differentially expressed, therefore, are likely of functional importance either for sexual or apomictic reproduction.

#### Cell cycle control involving protein degradation is likely of crucial importance during germline specification

Interestingly, based on homology or annotations, some of the DEGs are potentially involved in pathways recently identified to have regulatory roles during germline development and for restriction of reproductive fate (Kim et al., 2008; Gusti et al., 2009; Schallau et al., 2010; Schmidt et al., 2014; Singh et al., 2017; Tang et al., 2017; Zhao et al., 2017a; Zhao et al., 2017b; Cao et al., 2018).

For instance, 6 genes were among the identified candidates, which encode proteins containing F-box and/or associated interaction domains. Most of them were upregulated in apomictic nucelli. F-box proteins are involved in a multitude of biological processes in plants (Lechner et al., 2006; Stefanowicz et al., 2015). They are part of Skp1-Cullin1-F-box protein (SCF) ligase complexes, acting in the polyubiquitination-mediated 26S proteasomal degradation (Skowyra et al., 1997). As essential component of SCF complexes, they facilitate the regulation of many core cell cycle genes and are thus responsible for proper cell cycle progression including G1/S transition (Krek, 1998; Vodermaier, 2004). For example, *A. thaliana* SCF^SKP2A^ complex, containing the F-box protein SKP2A, positively regulates cell division (del Pozo et al., 2002; del Pozo et al., 2006; Jurado et al., 2008).

One of the F-box candidate genes, Bostr.15697s0319, even contains a cyclin-like and Skp2-like domain. Based on the homology of the interaction domains to SKP2, Bostr.15697s0319 is likely part of an SCF complex, guiding specific protein degradation during cell cycle progression. Interestingly, the *A. thaliana* SCF^FBL17^ complex, containing the F-box-like 17 (FBL17) protein, is described to act specifically in male germ cells, where it targets cyclin-dependent kinase inhibitors KIP-RELATED PROTEIN 6 (KRP6) and KRP7 to control germ cell proliferation (Kim et al., 2008; Gusti et al., 2009). KRPs in turn are also required to restrict female reproductive fate to only one MMC in *A. thaliana* ovules since it represses the cell cycle regulator RETINOBLASTOMA-RELATED 1 (RBR1) via CDKA;1 (Zhao et al., 2017b; Cao et al., 2018).

These interrelated findings demonstrate the essential function of F-box proteins for cell cycle regulation during germline development. Furthermore, similar to the male germline, they potentially are involved in specifying the female germline via SCF complex controlled regulation of KRPs (Kim et al., 2008). In line with this, F-box genes identified as differentially regulated, i.e. Bostr.15697s0319, might be part of SCF complex(es) governing cell cycle mechanisms that are distinct between apomictic and sexual germlines regarding apomeiosis and meiosis, respectively. Therefore, such KRP-dependent regulatory mechanisms or similar ones that regulate the cell cycle, might also play a role in *Boechera*.

Another differentially regulated gene, Bostr.25993s0534, encodes a TRAF-like family protein. TRAFs comprise a class of E3 ubiquitin ligases with characteristic RING finger, TRAF, and Meprine And TRAF Homology (MATH) domains, which enable them to mediate interactions between other TRAF members, receptors and several different intracellular signalling molecules (Ye et al., 1999; Zapata, 2003; Alvarez et al., 2010). In *A. thaliana*, the TRAF Mediated Gametogenesis Progression (TRAMGaP) is sharing the same interaction domains with Bostr.25993s0534 (Singh et al., 2017). It is an important regulator of plant germline development involved in restricting reproductive fate to a single MMC per ovule (Singh et al., 2017). Besides, the expression of several genes acting in germline specification and regulation of sporophyte to gametophyte transition was shown to be dependent on TRAMGaP. Importantly, these not only include RBR1 and the core meiotic gene *DYAD,* but also *AGO9, SGS3* and *RDR6* (Singh et al., 2017). While in *A. thaliana dyad* mutants largely lead to sterility, formation of triploid offspring fully retaining parental heterozygosity has been observed at low frequencies (Ravi et al., 2008). Furthermore, apospory-like phenotypes are described for *ago9*, *sgs3*, and *rdr6* mutants (Ravi et al., 2008; Olmedo-Monfil et al., 2010).

Based on the similarities of domains, it is tempting to speculate that Bostr.25993s0534 might have similar functions than TRAMGaP. However, future functional investigations is required to elucidate, if this gene is mediating apospory or diplospory by targeting *DYAD* or genes active in the small RNA pathway including *AGO9*.

Apart from this TRAF-like gene, additional DEGs higher expressed in the apomicts as in the sexual plants can be related to E3 ligases with described functions, i.e. Bostr.2983s0066 and Bostr.7867s0569. Bostr.2983s0066 is coding for an ubiquitin-like superfamily protein homologous to AT4G05310, for which a Cullin3A (CUL3A) dependent expression was described in *A. thaliana* (Dieterle et al., 2005). CUL3, which forms E3 ligase complexes with BTB and MATH domain proteins among others, is required for female gametogenesis and results in maternal effect embryo lethality in *cul3a cul3b* plants (Dieterle et al., 2005; Thomann et al., 2005). Bostr.7867s0569, encodes for a RING E3 ligase itself and is homologous to *ATRMA3*. Noteworthy, *ATRMA3* is expressed during pollen germination and pollen tube growth, and is co-expressed with *UBIQUITIN-CONJUGATING ENZYME 28 (UBC28)* (Wang et al., 2008). *A. thaliana* UBC28 in turn belongs to the UBC8 group of proteins known to interact with RING E3 ligases (Kraft et al., 2005; Stone et al., 2005). Interestingly, the *Boechera* homologue of *UBC28*, Bostr.29223s0097, was also identified to be significantly differentially expressed in this study, however it was higher expressed in the sexual accessions unlike the homologue of *ATRMA3*.

In the same line, HpARI7 completes the set of described RING E3 ligases potentially related to cell cycle regulation during megasporogenesis, since it is a candidate gene located on the apospory-linked *Hypericum APOSPORY* (*HAPPY*) locus in *Hypericum perforatum* (Schallau et al., 2010). The *HAPPY* locus in the *Hypericum* genome is in proximity of the homologous sequence of *ATHK1* (Schallau et al., 2010). Remarkably, the *Boechera* homologue of *ATHK1* was identified to be significantly downregulated in nucelli of all apomictic *Boechera* accessions in this study.

Taken together, a number of candidate genes identified in this study are coding for F-box proteins, E3 ligases or associated factors. Their *A. thaliana* homologues are directly or indirectly interrelated with previously described proteasomal degradation-mediated control of key cell cycle regulators and other targets, which influence reproductive fate decisions and germline development. This provides further evidence that tightly regulated protein degradation affecting cell cycle progression might be of crucial importance for governing the distinct specification and differentiation of apomictic and sexual germlines.

#### Involvement of stress responses and cell signalling in early megasporogenesis

Apart from cell cycle control involving protein degradation, other regulatory mechanisms, including stress response and hormonal pathways, have previously been described to be involved in megasporogenesis (Schmidt et al., 2015). Accordingly, some DEGs identified in this study are related to such pathways.

For example, the roles of oxidative stress and signalling including reactive oxygen species (ROS) are described to play roles in germline specification and possibly meiosis (Schmidt et al., 2015). In line with this, members of the *CYP450* superfamily of oxidoreductases are enriched in entire nucellar tissue of sexual *A. thaliana* compared to other tissues (Schmidt et al., 2011). Furthermore, they show cell cycle-dependent expression (Menges et al., 2002). Interestingly, recent findings demonstrate that also the *A. thaliana* cytochrome P450 gene *KLU* (*CYP78A5*) is involved in restricting female germline fate to only one MMC per ovule (Zhao et al., 2017a). Likewise, Bostr.7867s1594 is a homologue of *A. thaliana CYP81D4* and member of the *CYP450* superfamily, which potentially functions in response to ROS. It was significantly higher expressed in sexual than in apomictic nucelli. Thus, the detection of *CYP81D4* and other DEGs related to redox processes (Bostr.26833s0734 and Bostr.15774s0342) provide further evidence that regulation of redox homeostasis and response to oxidative stress, might contribute to the determination and differentiation of female sexual and apomictic germlines.

Beside its damaging properties, and thus triggering oxidative stress responses, ROS are also known to function as signalling cues between and within cells for example during male germline development (Kelliher and Walbot, 2012; De Storme and Geelen, 2014). Similarly, cell-to-cell signalling mechanisms are also required during female germline specification and differentiation (Grossniklaus and Schneitz, 1998; Koltunow and Grossniklaus, 2003; Schmidt et al., 2015; Zhou et al., 2017). In accordance with that, some DEGs can be attributed to cell signalling mechanisms, such as the homologue of *ATHK1*, which potentially acts as nonethylene phytohormone receptor (Tran et al., 2007). Other genes might be involved in auxin signalling within ovules, a relevant component of sporo- and gametogenesis in *A. thaliana* (Li et al., 2008; Pagnussat et al., 2009; Schmidt et al., 2011; Lituiev et al., 2013; Freire Rios et al., 2014; Schaller et al., 2015). These are namely Bostr.7867s1023, a transcription factor of the B3 family and Bostr.26833s0821, the homologue of *A. thaliana* GH3.6, for which auxin-responsiveness has been demonstrated previously (Nakazawa et al., 2001). Interestingly, AUXIN RESPONSE FACTORs belong to the same B3 super family of transcription factors and are well known key regulators of auxin signalling (Chandler, 2016).

In summary, putative functions of various identified candidate genes in apomictic and sexual *Boechera* nucelli are in line with the so far not fully understood complex regulatory network governing germline specification and differentiation in plants. Noteworthy, Tang et al. (2017) found DEGs with similar functional annotations in a comparative transcriptome analysis on entire flower buds of sexual and apomictic *Boehmeria tricuspis*. This further suggests that herein identified DEGs might be relevant for distinct apomictic and sexual fate decisions by facilitating oxidative stress response, auxin-driven cell-signalling and especially ubiquitination-mediated cell cycle control.

### Commonly and differentially regulated genes in apomictic and sexual nucelli

In accordance with the hypothesis that apomixis might be caused by asynchronous regulation of the originally sexual reproductive pathway, heterochronically shifted expression patterns in sexual and apomictic ovules of *Boechera* during megasporogenesis have already been reported (Sharbel et al., 2009; Sharbel et al., 2010). In the same line, the observed variation of expression patterns between nucellus tissues of similar developmental stages of the analysed apomicts might also relate to these genomic effects. It can be hypothesized that global deregulations of gene expression varies due to the different hybrid origins of apomixis in *Boechera* (Sharbel and Mitchell-Olds, 2001; Koch et al., 2003; Dobes et al., 2004a; Dobes et al., 2004b; Kiefer et al., 2009; Kiefer and Koch, 2012; Lovell et al., 2013). Furthermore, recent studies supported higher rates of mutation accumulation in apomictic as compared to sexual *Boechera* (Lovell et al., 2017) Still, if broad deregulations of gene expression due to genomic effects are causative or only correlated with apomixis is to date not fully understood. The rather small number of DEGs identified here, however suggests a small number of genes to be sufficient to mediate the switch from sexual reproduction to apomixis. This is in agreement with apomixis being genetically linked to only few loci in most species (Barcaccia and Albertini, 2013).

Apart from variations and DEGs observed, almost 40% of all genes in *Boechera* shared expression in all samples analysed, regardless of sexual or apomictic reproduction. These genes likely comprise common regulators of reproductive development during megasporogenesis, as similar tissues of related species were profiled. Generally speaking, it can be expected that the majority of expressed genes in a given organ or tissue type and at a certain developmental stage, e.g. the nucelli of ovules at megasporogenesis, are rather similar in closely related species. Moreover, exclusive gene expression for either reproductive mode appeared imbalanced, with more genes being expressed commonly in apomicts than in sexual nucelli. But due to lowest input of 38 dissected nucelli, less genes were identified to be active in sample LTM_2, compared to the other samples from sexual accessions. Thus, this apparent difference is probably not of biological relevance. Overall, the comparison of transcriptomes of entire *Boechera* nucelli with the transcriptomes of MMC or nucellus tissue of *A. thaliana* and the *B. gunnisoniana* AIC obtained from Schmidt et al. (2011 and 2014), indicated good accuracy of the data obtained.

In conclusion, in depth comparative analysis identified 45 genes, which were differentially expressed between sexual and apomictic accessions. Most interestingly, the newly identified genes can be related to regulation of genes and pathways previously described to relate to regulation of apomixis. The identification of these genes will provide a very good basis for future functional investigations. Thus, the study presented contributes to an understanding of the regulatory mechanisms governing apomixis and mediating specification of the female germline in apomictic and sexually reproducing plants in general.

## MATERIAL AND METHODS

### Plant material

Seeds of various accessions of the genus *Boechera* were kindly provided by Timothy F. Sharbel (Global Institute for Food Security, University of Saskatchewan, Canada) and Thomas Mitchell-Olds (Duke University, USA), or were transferred from the laboratory of Ueli Grossniklaus (University of Zürich, Switzerland). They were stratified in darkness for at least one week at 4^°^C, before surface sterilization and germination as described previously by Wuest et al. (2010). 1 – 2 weeks after germination plants were transferred to a mixture of soil (Einheitserde Classic CL ED73, Einheitserde Werkverband e.V., Sinntal-Altengronau, Germany) and sand (5:1), supplemented with Osmocote Exact Protect 5-6M fertilizer (ICL Specialty Fertilizers, Waardenburg, Netherlands). Plants were germinated and grown in a growth chamber with 55% relative humidity and in 16 h light / 8 h darkness at 20 – 22^°^C.

### Preparation of plant material for laser-assisted microdissection

For LAM, flower buds around the onset of megasporogenesis with ovules harbouring either a MMC or AIC (predominantly before meiosis or apomeiosis, respectively) were harvested. Tissue fixation and manual embedding was done as described by Wuest et al. (2010) with following modifications: Xylene gradient was applied to the tissue (25%, 50%, 75%, twice 100% xylene in EtOH, each step 1 – 1.5 hours at RT). Prior to blocking in Paraplast X-tra (Leica Biosystems, Nussloch, Germany), flower buds were incubated first overnight, then twice for 2 – 3 h in fresh Paraplast X-tra at 59 ^°^C. Embedded samples were stored at 4 ^°^C until further use. Thin sections of 7 μm were prepared from embedded flower buds with a Leica RM2255 microtome (Leica Biosystems, Nussloch, Germany), mounted on metal frame slides with 1.4 μm PET-membrane (Leica Microsystems, Wetzlar, Germany) and de-waxed according to Schmidt et al. (2011).

### Laser-assisted microdissection

LAM was performed with a Nikon Eclipse Ti microscope (Nikon Instruments Europe, Amsterdam, Netherlands) equipped with a mmiCellCut instrument (Molecular Machines & Industries (MMI), Eching, Germany) and the associated mmiCellTools software (version 4.5 #400). Nucellus tissue was isolated from thin sections using a 40x SPF Ph2 ELWD objective. On average approx. 60 nucelli sections were collected per cap and day. A control sample for each batch was obtained by collecting approx. 10 ovary sections from the same slides as the nucelli sections. Both, nucelli sections and control samples were stored dry on the caps at −80^°^C until RNA extraction.

### RNA extraction and amplification

Total RNA was extracted from LAM samples using the PicoPure^TM^ RNA isolation kit (Thermo Fisher Scientific) following manufacturer instructions, except that 15 μl extraction buffer was used and extracts from same accessions were pooled to obtain a final input amount of 38 – 226 nucelli sections per sample (Table S1). DNA was digested on the columns with RNase-free DNase I (Qiagen, Hilden, Germany). RNA integrity of control samples was determined using Agilent 2100 Bioanalyzer RNA Pico Chips following Schmidt et al. (2011). Consistently, good and reproducible RNA integrity with approx. RIN 7 on average was achieved (Figure S1A). A SMARTseq v4 Ultra Low Input RNA Kit for Sequencing (Takara Bio USA, Mountain View, USA) was used for linear amplification of mRNA derived from LAM samples following the manufacturer’s instructions. By this, high sensitivity and specificity was achieved (Figure S1B, S1C). Amplified cDNA was purified using NEBNext Sample Purification Beads (New England Biolabs, Ipswich, USA), dissolved in nuclease-free H_2_O (Ambion, Thermo Fisher Scientific, Waltham, USA) and stored at −20^°^C until library preparation.

### RNA sequencing

Libraries for RNA-Seq were prepared with a Nextera XT DNA Library Prep Kit (Illumina, San Diego, USA) following the user manual and purified with NEBNext Sample Purification Beads. Concentration and fragment size distribution of libraries were determined with Qubit and Bioanalyzer High Sensitivity DNA assays, respectively. The final equimolar library pool (30 nmol each) was again tested for homogenous composition (Figure S1D).

RNA-Seq was executed on one lane of a flow cell on an Illumina NextSeq 500 platform (Illumina, San Diego, USA) by the Deep Sequencing Core Facility of the University of Heidelberg. The 75 bp single-end protocol was applied with 94 cycles (including index read). Original data files are deposited in the NCBI SRA database with the accession number: SRP159014.

### Data analysis

#### Pre-analysis and processing of raw reads

Overall quality of raw reads was assessed with FastQC version 0.11.2 before quality and adapter trimming was performed with cutadapt version 1.14 (Andrews, 2010; Martin, 2011). Parameters were set to hard trimming of final base, NextSeq-specific quality trimming at 3’ end with phred score threshold 20, sequences of 5’ and 3’ adapters were provided to cutadapt as fasta files. Trimmed reads shorter than 30 bases were filtered out and remaining trimmed reads were again controlled with FastQC.

The *Boechera stricta* genome assembly from Lee et al. (2017) (Bstricta_278_v1.fa; obtained from the U.S. Department of Energy Joint Genome Institute on https://phytozome.jgi.doe.gov/pz/portal.html) was used for read mapping together with the corresponding gene annotation (Bstricta_278_v1.2.gene_exons.gff3), which was tidied with GenomeTools version 1.5.9 first (Gremme et al., 2013). Before mapping. the reference genome was indexed with standard settings using STAR version 2.5.3a_modified (Dobin et al., 2013). Mapped reads were sorted by scaffold position of reads and indexed using samtools version 1.6 (Li et al., 2009). The number of reads mapped to exons of each gene was determined with featureCounts from the Rsubread package version 1.20.6 (Liao et al., 2014) based on a published script (Schmid, 2017).

As quality controls, mapping statistics and gene body coverage were assessed with the modules bam_stat.py and geneBody_coverage.py from the RseQC package version 2.6.4 (Wang et al., 2012b). Duplication rates of reads were analysed with the package dupRadar version 1.0.0 of the Bioconductor software implemented in R, by plotting the expression density of three representative samples (covering the range of input amount; LTM_2, B12.1524, B12.1578_2; Figure S5) (Sayols et al., 2016).

#### Differential gene expression analysis

Read counts derived from featureCounts were subsequently used to analyse differentially expressed genes with the Bioconductor package edgeR version 3.12.1 implemented in R (Robinson et al., 2010). Samples were grouped by accession and reproductive mode, low expressed genes were filtered out (threshold defined as > 1 counts per million bases (CPM) in ≥ 2 libraries) and normalization factors for each library were calculated with the TMM-method (trimmed mean of M-values; Robinson and Oshlack (2010)). For statistical analysis of normalized read counts, common dispersion values were defined either as square of a set, constant biological coefficient of variation (bcv = 0.8) as previously used by Schmidt et al. (2014), if pairwise comparison were performed between individual apomictic and sexual accessions. Or it was estimated by edgeR as well as tagwise dispersion values (dispersion = 1.48), if samples of same reproductive modes were treated as biological replicates. Pairwise comparisons between sample groups were perform with exact test and Benjamini-Hochberg adjustment of the false discovery rate (FDR). Differential expressed genes with adjusted *p* values ≤ 0.05 were extracted and used for detailed analysis. To identify genes, which are differentially expressed in all comparisons (all sexual accessions against all apomictic accessions), an intersection of individual gene lists was calculated in R.

#### Data validation

Newly generated datasets were compared with previously published cell- and tissue type-specific transcriptome analyses of megasporogenesis in sexual *A. thaliana* and the triploid apomict B. *gunnisoniana* (Schmidt et al., 2011; Schmidt et al., 2014). Comparisons were restricted to commonly annotated *A. thaliana* homologues because of varying usage of reference genomes (*A. thaliana, B. stricta*) or transcriptomes (*B. gunnisoniana*). Genes were defined to be expressed, if they had either ≥ 10 mapped reads or present calls in ≥ 3 of 4 of microarray replicates (Schmidt et al., 2011; Schmidt et al., 2014; this study).

#### Data visualization

Overlapping gene expression was visualized as Venn diagrams using either the online tool Venny 2.1 (http://bioinfogp.cnb.csic.es/tools/venny/index.html; Oliveros, 2007 - 2015) or Biovenn (http://www.biovenn.nl/; Hulsen et al., 2008) for comparison of four or less datasets, respectively.

Heatmaps showing expression levels of differentially expressed genes were generated using the R package gplots version 3.5.0 (Warnes et al., 2016). Heatmaps were based on log2 transformed, TMM normalized read counts generated by featureCounts and NOISeq, and on applying hierarchal agglomerative clustering to the data as well as row scaling (Tarazona et al., 2011; Liao et al., 2014).

## ACKNOWLEDGEMENT

We thank Timothy Sharbel (Global Institute for Food Security, Canada), Thomas Mitchell-Olds (Duke University, USA) and Ueli Grossniklaus (University of Zurich, Switzerland) for providing *Boechera* seeds, Marcus A. Koch, Joachim A. Spatz (Max Planck Institute for Medical Research, Germany) and Jan Lohmann (Centre for Organismal Studies Heidelberg, Germany) for contributing necessary instruments and facilities, Markus Kiefer (Centre for Organismal Studies Heidelberg, Germany) for bioinformatics support, and Marc W. Schmid (MWSchmid GmbH, Switzerland) for helpful discussions and critical comments on the manuscript. We also acknowledge DFG for funding (SCHM2448/2-1) to AS.

**Author contribution**
AS conceived the project. AS and LZ planned the experiments. LZ conducted the experiments with support of DI and AS. LZ conducted the data analysis. LZ, AS, and DI interpreted the data. LZ and AS wrote the manuscript. All authors approved the manuscript.

